# 3D tree dimensionality assessment using photogrammetry and small unmanned aerial vehicles

**DOI:** 10.1101/023259

**Authors:** Demetrios Gatziolis, Jean F. Lienard, Andre Vogs, Nikolay S. Strigul

## Abstract

Detailed, precise, three-dimensional (3D) representations of individual trees are a prerequisite for an accurate assessment of tree competition, growth, and morphological plasticity. Until recently, our ability to measure the dimensionality, spatial arrangement, shape of trees, and shape of tree components with precision has been constrained by technological and logistical limitations and cost. Traditional methods of forest biometrics provide only partial measurements and are labor intensive. Active remote technologies such as LiDAR operated from airborne platforms provide only partial crown reconstructions. The use of terrestrial LiDAR is laborious, has portability limitations and high cost. In this work we capitalized on recent improvements in the capabilities and availability of small unmanned aerial vehicles (UAVs), light and inexpensive cameras, and developed an affordable method for obtaining precise and comprehensive 3D models of trees and small groups of trees. The method employs slow-moving UAVs that acquire images along predefined trajectories near and around targeted trees, and computer vision-based approaches that process the images to obtain detailed tree reconstructions. After we confirmed the potential of the methodology via simulation we evaluated several UAV platforms, strategies for image acquisition, and image processing algorithms. We present an original, step-by-step workflow which utilizes open source programs and original software. We anticipate that future development and applications of our method will improve our understanding of forest self-organization emerging from the competition among trees, and will lead to a refined generation of individual-tree-based forest models.

## 1. Introduction

Understanding how macroscopic patterns of forests emerge as a result of self-organization of individual plants and how ecosystems respond to environmental gradients and disturbances that occur at different spatial and temporal scales has long been reported as a largely unresolved fundamental ecological challenge (Levin, 1998). The phenotypic plasticity of individual trees is regarded as the major biological determinant of self-organization, structure, and dynamics of forested ecosystems and their response to natural and anthropogenic disturbances (Strigul, et al., 2008; Strigul, 2012). Unique patterns of tree plasticity have been identified across ecological and species groups, for instance, in conifers (Loehle, 1986; Umeki, 1995; Stoll & Schmid, 1998) and broad-leaf trees (Woods & Shanks, 1959; Brisson, 2001); and biomes, including tropical (Young & Hubbell, 1991) and temperate ecosystems (Gysel, 1951; Frelich & Martin, 1988; Webster & Lorimer, 2005). Failures to predict growth at the individual tree level with acceptable accuracy have been attributed to the heterogeneity in geomorphic and climatic phenomena affecting tree survival and growth, but primarily to inadequate information on the size, shape, and spatial distribution of interacting trees (Strigul, 2012).

National Forest Inventory (NFI) systems are a major source of systematic, spatially distributed, and repeated individual tree measurements obtained during field visits of established plots. A review of NFI field protocols and data quality standards reveals that very precise measurements are prescribed for tree stem diameter at breast height, and for fixed-area field plots, distances used in determining whether a tree stem center is within the plot area. Where recorded, the relative position of tree stems within a plot and tree height is measured accurately. Some vegetation parameters such as shrub and forb percent cover, crown base height, and crown compaction ratio are assessed ocularly, and therefore should be regarded more as estimates rather than measurements. Owing to cost, complexity, and logistic constraints such as visibility, crown width and other specialized tree dimensionality measurements are obtained only during special projects.

Information on individual trees over large areas is feasible only via processing of remotely sensed data. High (submeter) resolution space-or airborne spectral imagery has been used to identify and delineate individual tree crowns (Wulder et al., 2000; Leckie et al., 2005; Hirschmugl et al., 2007; Skurikhin et al., 2013), and to assess parameters of crown morphology such as height, radius, and surface curvature (Gong et al., 2002; Song, 2007) using various modeling approaches. Information extracted by manual interpretation of aerial photographs has often been used as surrogate of field measurements for model development and validation (Gong et al., 2002; Coulston et al., 2012). The advent of Light Detection and Ranging (LiDAR) technology enabled 3D measurements of vegetation over forested landscapes. Operated mainly from airborne platforms, LiDAR instruments emit short pulses of light that propagate through the atmosphere as a beam of photons and are backscattered to the instrument from illuminated targets. The loci of interactions with objects or object parts along a beam’s trajectory are determined with decimeter precision and reported as points georeferenced in three dimensions. The collection of points generated across all pulses is referred to as a point cloud. A typical LiDAR data set of a forested scene comprises points from the entire volume of tree crowns and ground surfaces. Models operating on metrics that describe the spatial distribution of above-ground points have been proven useful for assessing area-based forest inventory parameters such as wood volume and biomass (Zhao et al., 2009; Sheridan et al., 2015). With high-density LiDAR data, a single mature tree can be represented by many, up to hundreds of points, conditions conducive to a precise assessment of its dimensions, including height and crown width (Popescu et al., 2003; Andersen et al., 2004). Often however, the token representation of lower canopy components and ground surfaces in LiDAR data sets caused by substantial attenuation of pulse energy in dense, multistory stands leads to less accurate estimates of tree dimensionality (Gatziolis et al., 2010; Korpela et al., 2012). Terrestrial LiDAR systems operated from ground or near-ground locations deliver point cloud densities orders of magnitude higher than those generated by using airborne instruments, enabling detailed and precise reconstructions of individual trees (Côté et al., 2009). Modeling of crown morphology supported by terrestrial LiDAR data has been shown effective in assessing how trees grow in response to competition between and within crowns (Metz et al., 2013). Point clouds generated from single scanning locations always contain gaps due to partial target occlusion, either from parts of the targeted tree itself or from surrounding vegetation. As occlusion rates, gap frequency, and gap size increase with canopy height, the error levels in tree dimensionality estimates obtained by processing these point clouds also increase with height (Henning & Radtke, 2006; Maas et al., 2008). Ensuring that estimate precision meets established standards necessitates scanning targeted trees from multiple locations and then fusing the individual point clouds, a complication that often is logistically complex and costly.

To date, precise tree crown dimensionality and location data supportive of a rigorous modeling of individual tree growth has been inhibited by feasibility, logistics, and cost. Measuring crown characteristics by using established inventory methods is very time consuming and hardly affordable outside special projects. Existing remote sensing methods of measuring tree crowns provide only partial crown reconstructions. Airborne LiDAR data acquisitions require prolonged planning and are costly. As an example, the minimum cost for a single airborne LiDAR acquisition with common specifications in the US Pacific Northwest exceeds $20,000 irrespectively of acquisition area size (Erdody & Moskal, 2010). Transferring to and operation of terrestrial LiDAR instruments in remote forest locations and challenging terrain is both labor intensive and time consuming. As a result, the assessment of tree growth and competition relies on numerous simplifying, albeit often unjustified, assumptions such as of trees with symmetric, vertical, perfectly geometric crowns growing on flat terrain, and illuminated by omnidirectional sunlight. These assumptions propagate through modeling efforts and ultimately reduce the validity of model predictions, thereby decreasing their utility (Munro, 1974; Strigul, 2012).

Recently, unmanned aerial vehicles (UAVs) equipped with inexpensive, off-the-shelf panchromatic cameras have emerged as a flexible, economic alternative data source that supports the retrieval of tree dimensionality and location information. Flying at low altitude above the trees and with the camera oriented at a nadir view, UAVs acquire high-resolution images with a high degree of spatial overlap. In such conditions, a point on the surface of a tree crown or a small object on exposed ground is visible from many positions along the UAV trajectory and is depicted in multiple images. Automated photogrammetric systems based on computer Vision Structure from Motion (VSfM) algorithms (Snavely et al., 2008) explore this redundancy to retrieve the camera location the moment an image was acquired, calculate an orthographic rendition of each original image, and ultimately produce a precise 3D point cloud that represents objects (Dandois & Ellis, 2010; Rosnell & Honkavaara, 2012). Application of VSfM techniques on UAV imagery has enabled accurate 3D modeling of manmade structures, bare ground features, and forest canopies (de Matías et al., 2009; Danbois & Ellis, 2013; Dey et al., 2012). Automated image processing is now supported by open-source and commercial software packages.

Image acquisitions with nadir-oriented cameras onboard UAVs, however, face the same issues as airborne imagery; the great majority of points in derived clouds are positioned near or at the very top of tree crowns. The representation of crown sides tends to be sparse and contains sizeable gaps, especially lower in the crown, a potentially serious limitation in efforts to quantify lateral crown competition for space and resources, as in the periphery of canopy openings. In this study, we extend UAV-based image acquisition configurations to include oblique and horizontal camera views and UAV trajectories around trees or tree groups at variable above-ground heights to achieve comprehensive, gap-free representations of trees. To overcome the challenges imposed by these alternative UAV/camera configurations, we evaluated many UAV platforms and open-source VSfM software options, and developed original, supplementary programs. To determine whether comprehensive tree representations are attainable, we initially processed synthetic imagery obtained via simulation. We finally evaluated the efficacy and performance of our workflow targeting trees of different species, shapes, sizes, and structural complexity.

## 2. Method development and testing

### 2.1. Image processing

The procedure that uses a set of images exhibiting substantial spatial overlap to obtain a point cloud representing the objects present in the images contains three main steps: feature detection, bundle adjustment, and dense reconstruction. To implement this procedure, we have carefully examined a variety of software available for image processing. The workflow presented below was found by experimentation to be the most efficient for our project. We employed a sequence of computer programs, most of which are available as freeware or provide free licenses to academic institutions. The software used includes OpenCV libraries, VisualSFM, CMVS, SURE, OpenGL, and Mission Planner, with each of them accompanied by a comprehensive manual. Considering that the majority of the software listed above evolves rapidly, we intentionally refrained from duplicating here elements of associated manuals to which we refer a reader in addition to our presentation.

Feature detection is based on the identification of image regions, often called keypoints, pertaining to structural scene elements. Thanks to image overlap, these elements are present in multiple images, but because their position relative to the focal point of the camera is image-specific, they are depicted in different scale and orientation (Figure 1). Illumination differences and image resolution can impose additional feature distortions. Algorithms used in feature detection explore principles of the scale-space theory (Lindeberg, 1998). According to this theory, a high-resolution image can be perceived as a collection of scene representations, called octaves, in Gaussian scale space. The scale space can be obtained by progressively smoothing the high-resolution image, an operation analogous to a gradual reduction of its resolution. If robust against changes in scale and orientation, the characteristics of a keypoint identified on a given octave of one image can be used to identify the same keypoint on other images. The algorithms proposed for feature detection in this context include the Scale Invariant Feature Transform (SIFT) (Lowe, 2004), the Speeded Up Robust Features (SURF) (Bay et al., 2008), and the Oriented FAST and Rotated BRIEF (ORB) (Rublee et al., 2011). We employed SIFT in our workflow as it is currently the reference approach in the field of computer vision. To identify keypoints, SIFT initially applies to each image octave an approximation of the Laplacian of Gaussian filter known as Difference of Gaussians, an efficient edge detector. Identified SIFT keypoints are circular image regions, each described by a set of parameters: the image coordinates at the center of the region, the radius of the region and an angle. The radius and angle of each keypoint serve as scale and orientation indicators respectively (Figure 1). Keypoints are further characterized by a descriptor of their neighborhood, determined from the values of pixels in the vicinity of the keypoint’s center and usually encoded into a vector of 128 values. By searching for keypoints at multiple scales and positions, SIFT is invariant to image translation, rotation, and rescaling, and partially invariant to affine distortion and illumination changes. It can robustly identify scene features even in images containing substantial amounts of noise or under partial occlusion.

**Figure 1.**
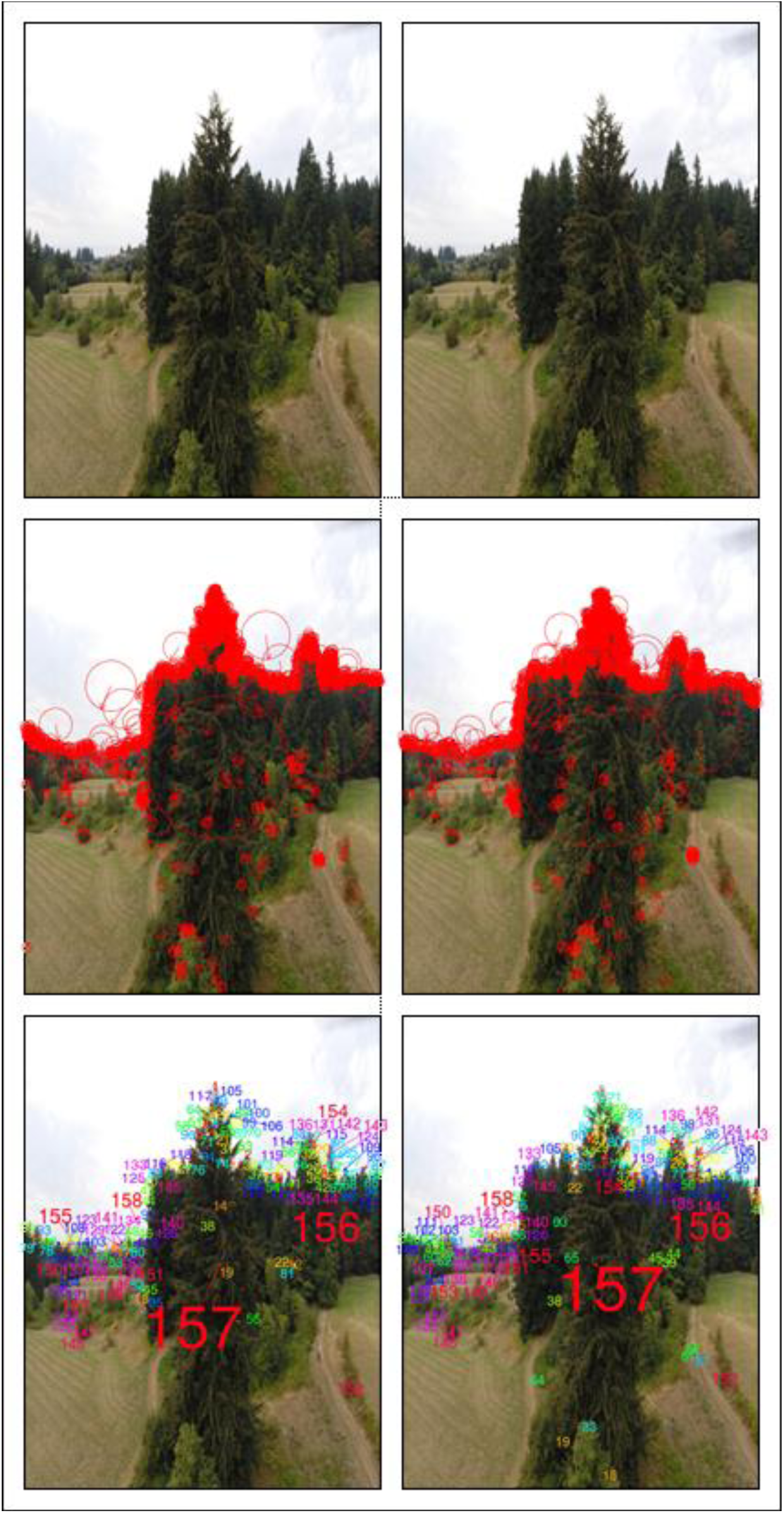
SIFT-based scene keypoint detection and matching on two overlapping images. Top: Original images; Middle: 1464 (left) and 1477 (right) keypoints with arrows denoting orientation and radii scale; Bottom: 157 keypoint pairs, matched by color and number.

The bundle adjustment process initially compares keypoint descriptors identified across images to determine two similar images. Then, an optimization procedure is performed to infer the positions of cameras for these two images. Remaining images are added one at a time with relative positions further adjusted, until camera locations become available for all images. The optimization often uses the Levenberg-Marquardt algorithm (Levenberg, 1944; Marquardt, 1963), a general purpose non-linear optimization procedure. Heuristics and prior information, such as GPS coordinates of UAV locations at the moment an image is acquired, can be included to improve convergence speed. In the end, the spatial positions and orientations of all cameras are triangulated using the keypoints identified in the previous step. At the conclusion of bundle adjustment a so-called sparse 3D model that contains the 3D positions of all identified features becomes available. We implemented the feature detection and bundle adjustment components of our workflow in VSFM software (Wu, 2013; Wu et al, 2013).

In dense reconstruction, the final processing step, all image pixels, not only keypoints, along with the positions and orientations of each camera, are merged into a single high-density structure. This is achieved by matching pixels with similar value across pictures with respect to the epipolar geometry constraints (Zhang et al., 1995) of the sparse model. The epipolar geometry is defined for each image pair. It includes a baseline connecting the locations of the two cameras that are known from the sparse model, the oriented image planes, the image locations where image plane and baseline intersect known as epipoles, and the epipolar lines connecting a camera location with a pixel on the image plane. By restricting searches for a pixel match along the epipolar lines, processing is greatly expedited. In our workflow, we considered CMVS (Furukawa & Ponce, 2010) and SURE (Rothermel et al., 2012), two state-of-the-art, freely available multi-core implementations, which adopt different strategies to generating the dense model. CMVS is a patch-based method which starts from matched keypoints and generates local models of object surfaces, or patches, in the immediate neighborhood of the keypoints. These patches are then expanded until their projections on the original pictures eventually form a dense tiling. SURE’s approach is based on the computation of depth maps for a set of reference images, based on the disparity between these images and other images obtained from nearby, according to the sparse model, positions. Each depth map provides a dense model of pixels equivalent to a local reconstruction from one reference viewpoint. All partial reconstructions are eventually merged to obtain a dense reconstruction for the entire scene.

The sparse and dense reconstructions obtained from a set of overlapping images are configured in the same internal coordinate system and scale. Conversion to real-world orientation and coordinate system is a prerequisite for meaningful measurements of reconstructed objects or for comparisons with ancillary spatial data. Such conversions can be performed manually on the reconstructed scene, assuming reference *in-situ* measurements of object dimensionality are available. In this study, we used an alternative, automated approach. The latitude, longitude, and elevation of camera locations recorded by a recreational-grade GPS device onboard the UAV were converted to orthographic Universal Transverse Mercator (UTM) coordinates using a GDAL (2015) reprojection function. The rotation/translation matrix linking the UTM and sparse model coordinates of the camera positions was then calculated via maximum likelihood, and applied to convert the sparse model coordinates system to UTM. All subsequent processing by CMVS and SURE were performed on the UTM version of the sparse model.

### 2.1.1 Image calibration

All imaging systems introduce a variety of distortions onto acquired imagery. The magnitude of the distortion is usually negligible in professional systems, but it can be substantial for inexpensive, off-the-shelf cameras used in structure from motion applications (Balletti et al., 2014). Most software, including VSfM, perform internal image calibration using information on the focal length of the lens, usually stored in the header of the image, and a generic rectification process, or undistortion as it is commonly called. Departures between the actual distortion and the one anticipated by the generic rectification process reduce the spatial accuracy of reconstructed objects. Using simulated and UAV-based, nadir looking imagery featuring sparse and low vegetation on flat land, Wu (2014), the author of the VSfM software, documented that scene reconstructions obtained by using the generic image calibration model present in VSfM produced a macroscopically concave ground surface, an artifact attributed to imprecise image calibration. To avoid artifacts, we first calibrated all cameras used in this study with the efficient procedure described in the OpenCV image processing library (Bradski, 2000), and then instructed VSfM to skip the generic image calibration process. Separate calibrations were performed for each operating mode of each camera. As expected, and evident in Figure 2, calibration effects were more discernible near the periphery of the image. The convex scene horizon in the original image appears flat and horizontal after calibration and the local road pavement on the lower left part of the original image is excluded from the calibrated version.

**Figure 2.**
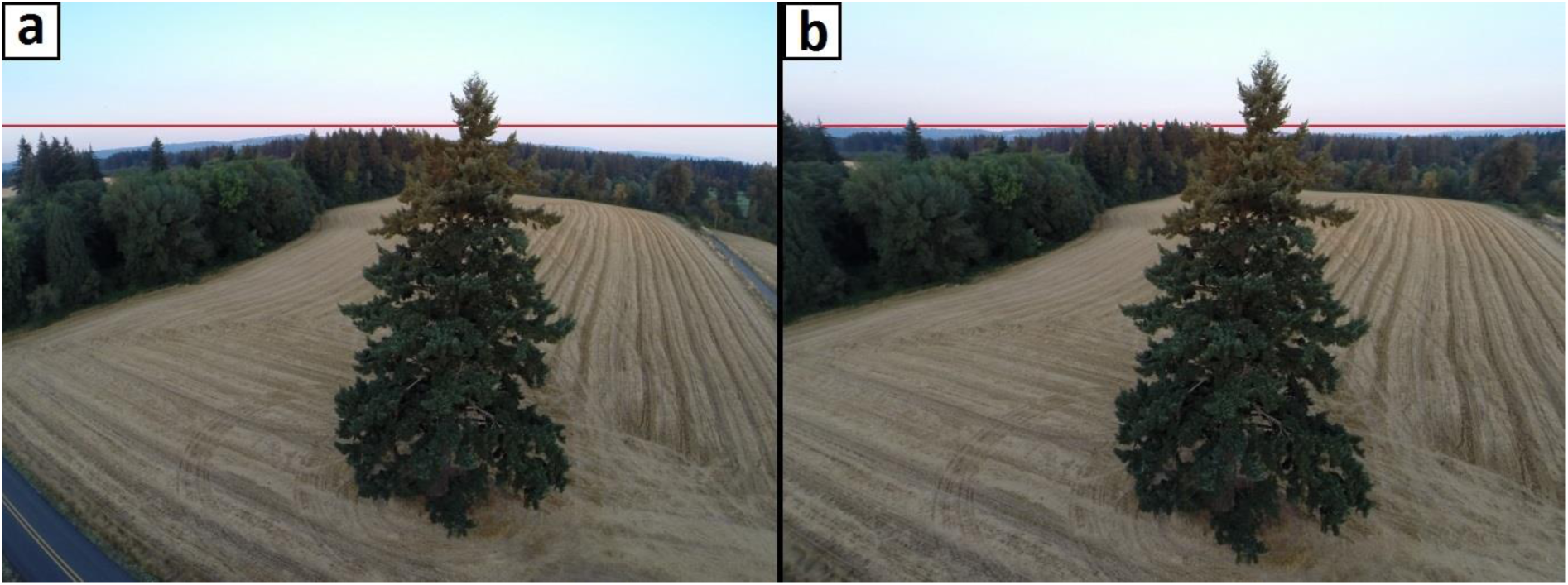
Removal of lens distortion. Demonstration of a. original, vs. b. OpenCV-calibrated lateral tree image obtained with a UAV-based GoPro camera at an above-ground altitude of 18 meters. Horizontal red line drawn to illustrate form of horizon in each version of the image.

### 2.2. Simulation-based assessment of image-based tree reconstruction accuracy

Upon initial consideration, the accurate and detailed reconstruction of objects characterized by complex structure and geometry, such as trees, using image-based techniques may be deemed an ill-fated effort. The main reason for pessimistic prognoses is that the aforementioned methods and algorithms used in processing the imagery anticipate planar surfaces as structural elements of the objects and well-defined edges at object surface intersections. Except for the lower part of the main stem of large trees, sizeable and homogeneous surfaces separated by crisp boundaries are absent in trees. A second reason is that trees are not opaque objects. Even in high foliage and branch density conditions, portions of scene background are clearly visible through the tree crowns. The see-through-crown phenomenon can be overlooked in nadir-oriented imagery where the forest floor is acting as tree background, but it is often rather pronounced in lateral imagery where the depth of the part of the scene situated behind the trees can be large. The term ‘lateral’ is used here to describe images acquired with the UAV positioned to the side of the tree and lower than the tree top. The effects of substantial differences in parallax between tree components and background depicted only pixels apart in lateral tree imagery, and high rates of component occlusion, are likely analogous to image distortion, a condition to which the SIFT algorithm is only partially invariant. Furthermore, the upper parts of tree crowns depicted in lateral imagery can have the sky as background instead of the typically darker vegetation or terrain background present in nadir-oriented imagery. Drastic changes in background brightness, for instance, from sky to vegetation and back to sky, behind a given part of a tree crown that appears across multiple overlapping lateral images, influence the red, green, and blue (RGB) values of image pixels corresponding to that crown part. The ensuing variability in pixel values often mimics effects induced by differences in diurnal solar illumination regimes. Illumination variability is another condition to which SIFT is only partially invariant. We used simulation and synthetic images to evaluate the robustness of our standard workflow to the idiosyncrasies of lateral tree imagery described above. We relied on terrestrial LiDAR data representing a collection of free-standing trees, each scanned from multiple near-ground locations. The scanning was performed in high-density mode with the laser beams distributed in fine horizontal and vertical angular increments (0.4 mrad). Each point in the generated clouds was furnished with RGB values extracted from panchromatic imagery captured by the LiDAR instrument during the scanning. Details on the data acquisition are available in Gatziolis et al. (2010). The RGB-colored point cloud of each tree was then visualized in an OpenGL interface (Shreiner, 2009) with perspective rendering (Figure 3a). In this virtual visualization environment, RGB-colored snapshots of each scene, henceforth referred to as synthetic images, can be obtained without limitations on image number, resolution, amount of spatial overlap, and format type. By specifying the trajectory, orientation, snapshot frequency, and field of view of the virtual camera and also the pixel dimensionality of the OpenGL interface, we can control the scale at which targeted trees, or parts of trees, are represented in the synthetic imagery. The background can be adjusted to resemble the overall scene illumination conditions effective during the acquisition of the terrestrial imagery, including illumination adjustments along azimuth and sun elevation angle gradients. Synthetic images generated by exercising combinations of these options yield very realistic approximations of imagery obtained onboard the UAVs, with the additional advantage that the dimensionality of the objects depicted in the imagery is precisely known. Point clouds generated by processing the synthetic imagery can then be compared to the original terrestrial LiDAR point cloud to evaluate the accuracy and precision of object reconstructions.

**Figure 3.**
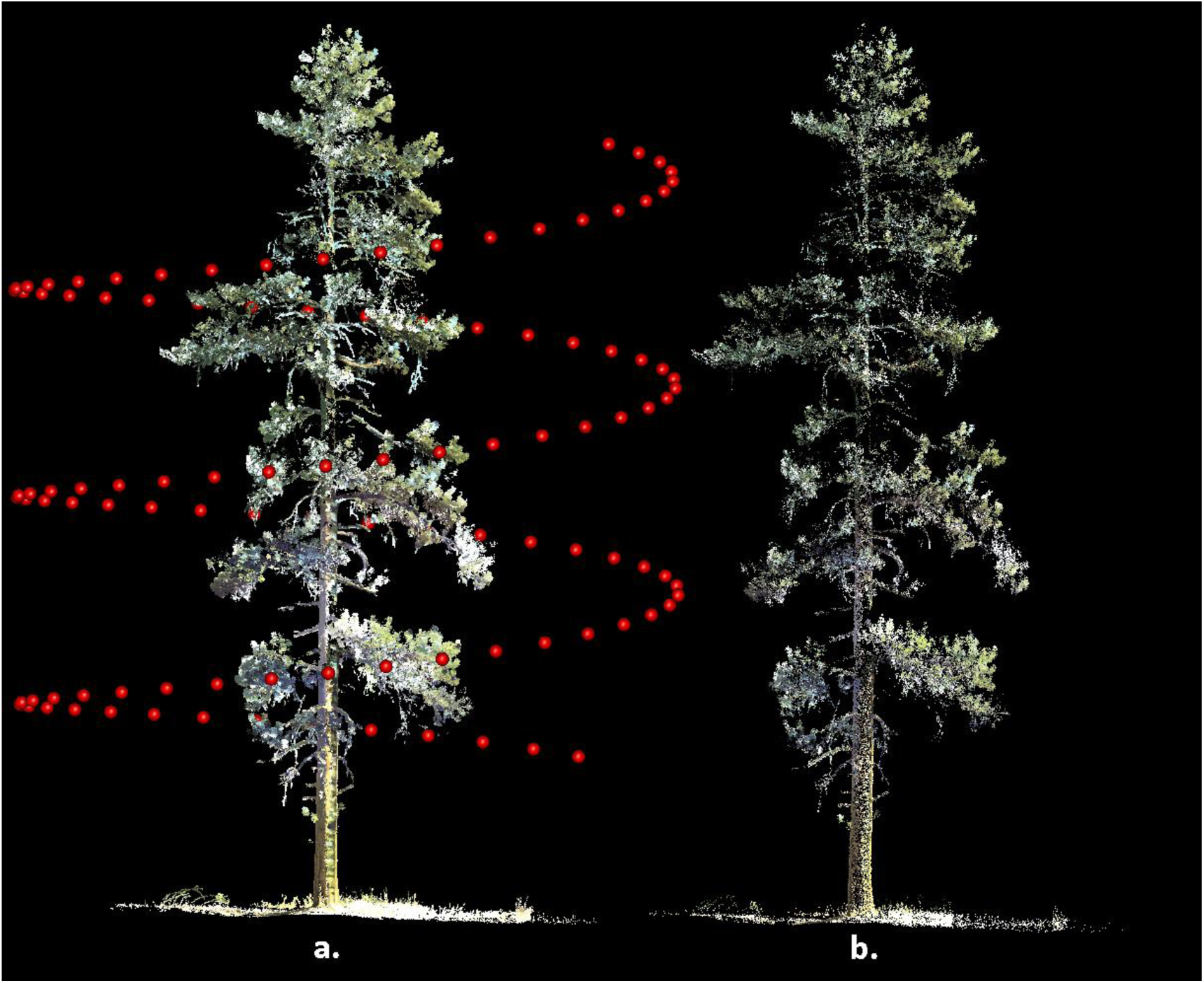
3D reconstruction in simulation. a. Perspective view of point cloud acquired with terrestrial LiDAR and camera locations (red spheres) used to obtain virtual images of the scene. b. Scene reconstruction obtained by processing of the images.

For our simulations we employed a 2500 by 2000 pixel (5 Mp) virtual camera. The camera was positioned on a circular trajectory centered on the crown of each of the trees depicted in the terrestrial LiDAR point clouds. The camera trajectory was either aligned to a horizontal plane elevated to approximately the vertical middle of the crown, or along a spiral ascent from the 15^th^ to the 85^th^ percentile of tree height (Figure 3a). Camera distance to the nearest part of a crown was between 10 and 15m and scene background was set to black. Between 100 and 250 synthetic images were acquired for each tree and trajectory combination, initially in BMP (bitmap) format and subsequently converted to the Joint Photographic Experts Group (JPEG) format, required by VSFM, using a maximum quality setting in ImageMagick, an open-source software suite (http://www.imagemagick.org). The synthetic imagery for each tree was processed with VSFM using standard settings, and the coordinates of the resulting point clouds generated at the sparse reconstruction stage were converted to the coordinate system of the terrestrial LiDAR data using the locations of the virtual camera known from the simulation settings. Dense reconstructions were obtained by using SURE with standard setting plus an option to ignore synthetic image regions with very low variability in pixel values, as those representing the scene background.

The original Terrestrial LiDAR and dense reconstruction point clouds for each tree were compared in voxel space (Popescu & Zhao, 2008; Gatziolis, 2012). In this setting, the bounding box of a point cloud is exhaustively partitioned into discrete, equally-sized cubical elements, called voxels. Those voxels containing one or more points are labeled ‘filled’, all others remain empty. By ensuring that the terrestrial and reconstruction voxel spaces have the same origin and voxel size, we were able to calculate the spatial correspondence of filled voxels between the two clouds and the rates of omission and commission, and identify parts of the voxel space where correspondence is better or worse than in other parts. The size, or resolution, of the voxels was set to 2cm, in response to the angular resolution of the terrestrial LiDAR beams at the mean distance between trees and LiDAR instrument.

### 2.3. UAV platform characteristics and image acquisition procedures

After a preliminary evaluation of several commercially available UAV platforms, we focused on an APM:Copter (http://copter.ardupilot.com), a hexacopter rotorcraft (Figure 4), because of its easily modifiable architecture and open source software for flight control. We also used a commercial IRIS quadcopter developed by 3DRobotics (http://3drobotics.com). The components of the customized hexacopter and their purchasing prices are shown in Table 1. Both systems feature gyroscopes and GPS receivers. Compared to systems available in the market, our hexacopter is an inexpensive but versatile configuration whose component acquisition cost is expected to drop substantially in the future as UAV technology evolves and its popularity continues to increase.

**Figure 4.**
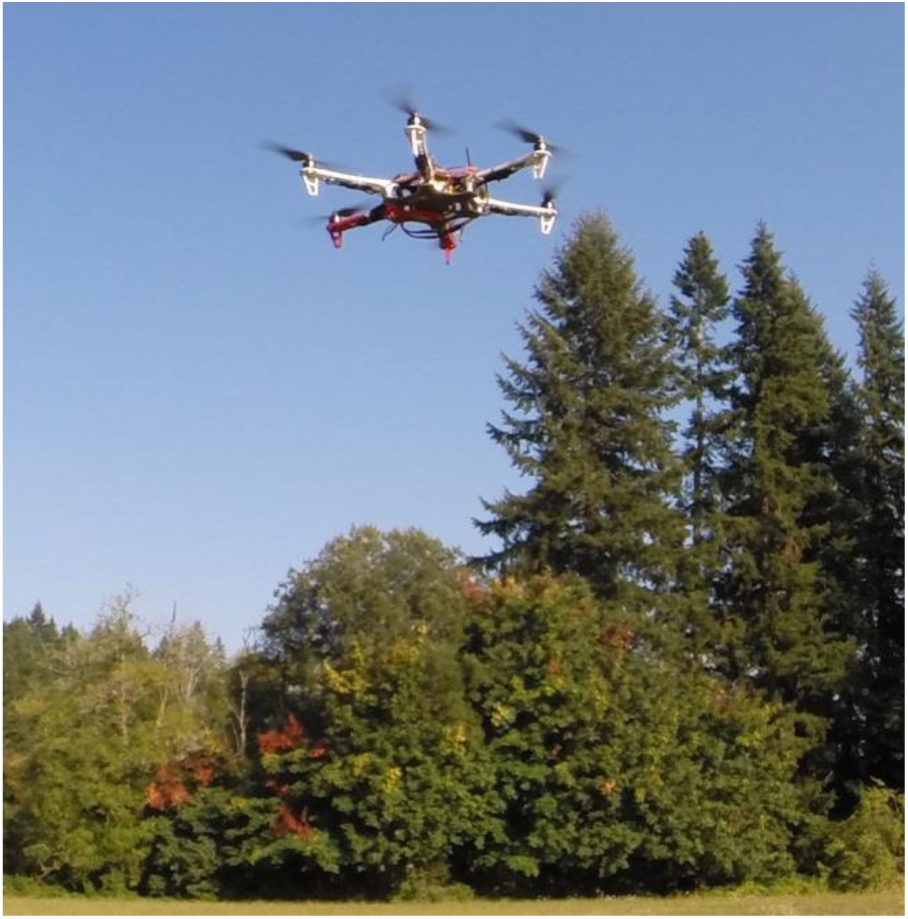
Custom built UAV hexacopter used to collect imagery data in this study.

**Table 1.**
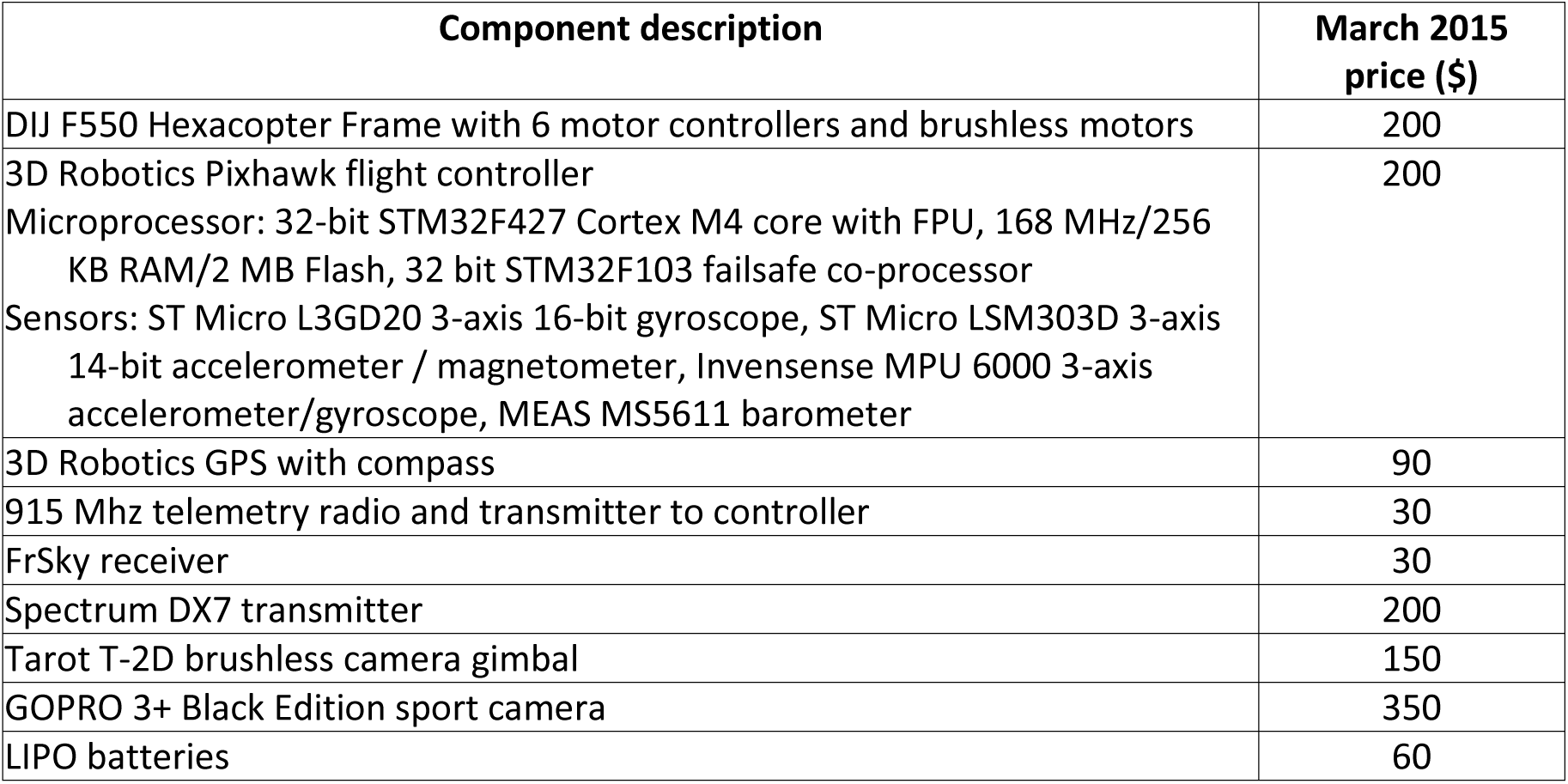
Specifications and prices of customized UAV platform used in this study at the time of writing.

Both UAVs used in this study can be operated either autonomously along a predefined trajectory or manually. The manual flight control requires expertise and continuous line of sight between the system and the operator. Maintaining nearly constant planar and vertical speed and orientation of the onboard camera towards the target is challenging, even for operators with years of experience. Experimentation confirmed that imagery acquired with manual flight control exhibits variable rates of overlap between frames captured sequentially. Smaller components of the targets are sometimes depicted in too few frames or are missing completely, while others appear in an excessive number of frames. For these reasons, it was decided to rely on autonomous flights configured by prior mission planning, and reserve the manual mode only for intervention in the event of an emergency.

#### 2.3.1. Characteristics of the imaging system

We conducted extended trials with several cameras, including the sport GOPRO 3+ Black Edition (http://gopro.com/), Ilook Walkera (http://www.walkera.com/en/) and Canon PowerShot (http://www.canon.com). The evaluations involved all operating modes offered by each camera, including normal, wide, and superwide zoom settings, as well as acquiring video and then extracting individual frames with post-processing. At the conclusion of the trials, we selected the GOPRO 3+ Black Edition operated in photography mode, and normal, 5 Mp resolution. Acquired frames were stored in JPEG format to the camera’s flash card. We rarely achieved event partial tree reconstruction using the alternative settings, likely because of the magnitude of distortion embedded into the imagery.

#### 2.3.2. Mission planning

The objective of the mission planning phase is to optimize the UAV trajectory, attitude, speed, and were applicable, the view angle of the camera gimbal for image acquisition. The gimbal is a hardware component which allows the orientation of the camera to be modified during the flight relative to the platform. Dynamic, trajectory-location-specific adjustments of camera orientation can be used to ensure that the target is centered on the images, especially when the UAV trajectory is not along a horizontal plane. During mission planning the image acquisition frequency is also considered. After rigorous evaluation of various UAV trajectory templates (Figure 5), we determined that the optimal reconstructions of trees are achieved when sequential images have a field-of-view overlap of approximately 70%. In this configuration, the nominal mean number of images where a part of a targeted tree would be present in is 3.4. Once determined, a trajectory template is centered on the target and scaled so that during the actual flight the mean camera-tree distance, platform speed, and image acquisition frequency will generate images exhibiting the targeted field-of-view overlap. The process is perceptually simple, but technically complex considering that all directional and attitudinal vectors of the UAV have to be converted to instructions passed to the UAV controller. Thankfully, it can be streamlined by using Mission Planner, an open-source software suite developed by Michael Osborne (http://planner.ardupilot.com). Mission Planner relies on user input and georeferenced imagery of the targeted area and tree(s), to establish the geographic (latitude and longitude) coordinates of the UAV’s starting and ending position and trajectory. A small set of high-level Mission Planner commands can accomplish even complex trajectory templates. All templates shown in Figure 5 require only 5 commands (Table 2). Our typical setup uses a location positioned in the middle of an open area for both the start and end of the flight. The UAV would initially ascend vertically above its starting location to a pre-specified height, then move horizontally to the beginning of the trajectory, complete it, and finally return to the starting location. In the present development state of our system, it is the user’s responsibility to ensure that the designed flight path is free of other objects, an easy to achieve requirement considering the wealth of georeferenced, high resolution, publicly available aerial photographs (Figure 6). The Mission Planer is also used to convert telemetry data of camera locations the moment images were acquired, provided by the GPS receiver stored to the onboard flash memory card, to an accessible format. As detailed in Section 2.1, these locations are later paired to those calculated during the sparse reconstruction processing phase to adjust the scale and georeference of reconstructed objects.

**Figure 5.**
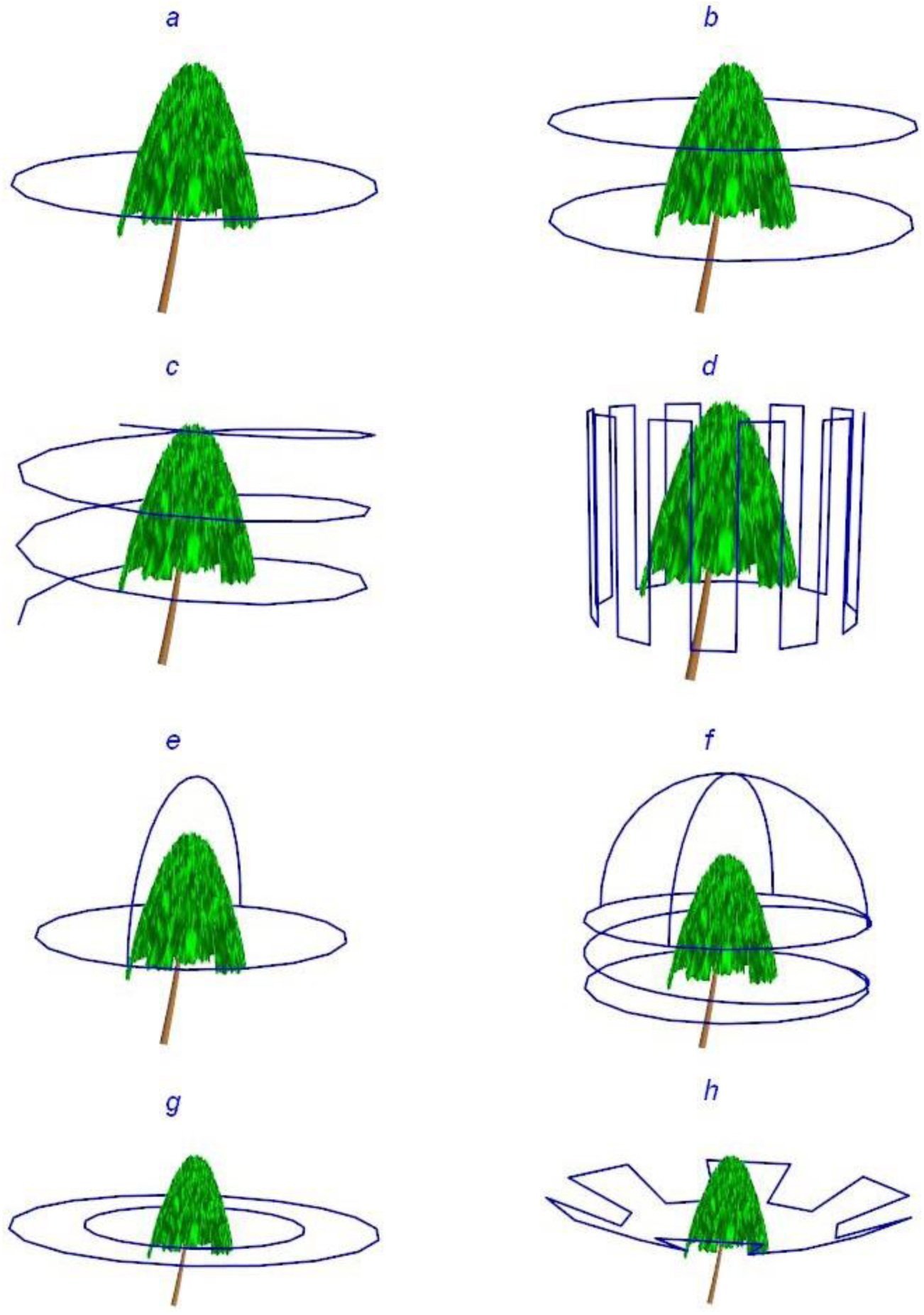
Different UAV trajectories tested for image acquisition. a. circular, at constant height; b. ‘stacked circles’, each at different above-ground height, for tall trees (height more than 20 m); c. spiral, for trees with complex geometry; d. vertical meandering, targeting tree sectors; e. clover, for trees with wide, ellipsoidal tree crowns; f. ‘spring-hemisphere’, designed for trees with flat-top, asymmetrical crowns; g. ‘nested circles’, centered on the tree; and h.’jagged saucer’, designed for trees with dense foliage but low crown compaction ratio.

**Figure 6.**
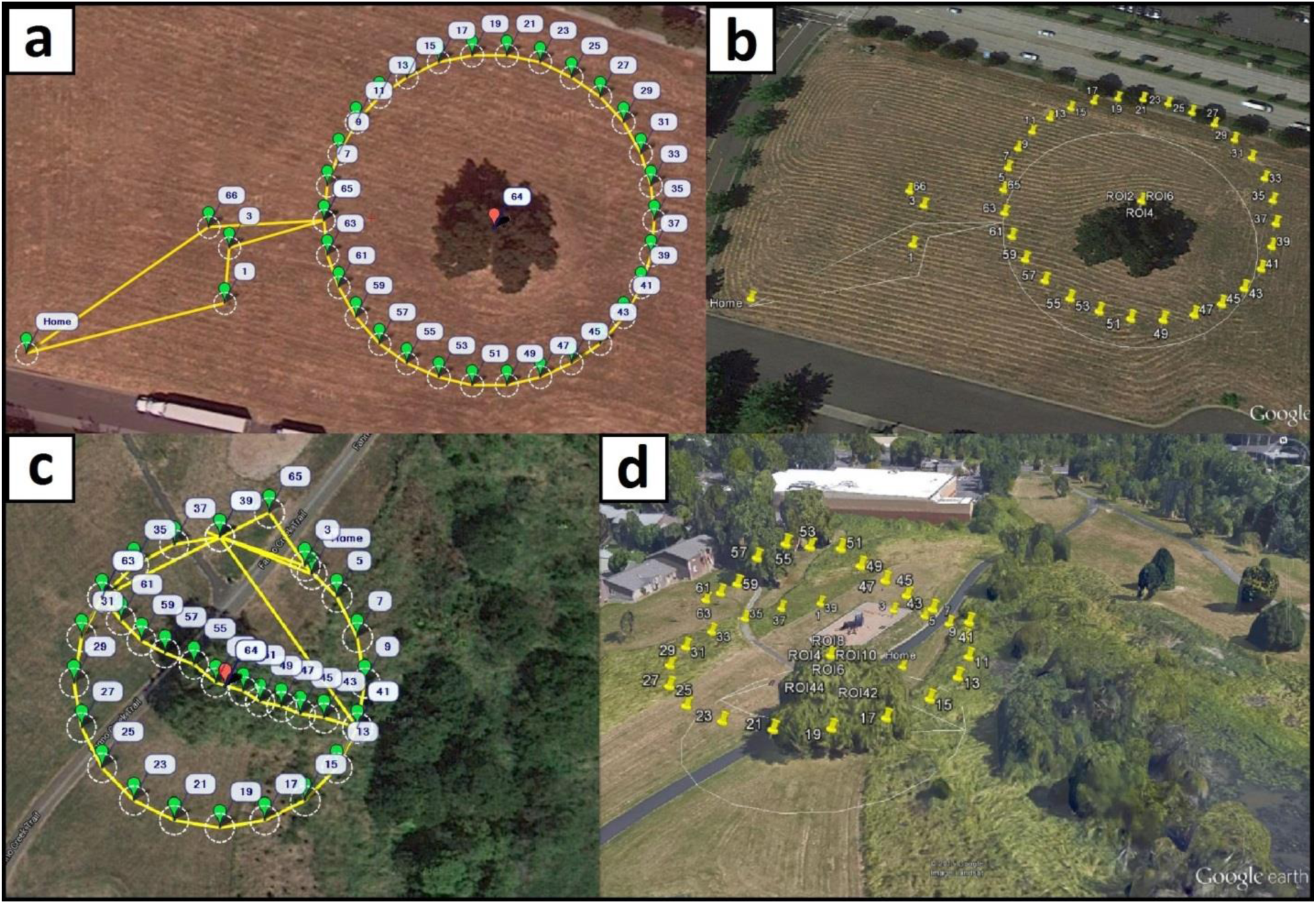
Visualization of designed and accomplished UAV trajectories. a. and c. circular and clover templates as seen in Mission Planner with yellow lines showing the flight paths, green balloons indicating waypoints, and red balloons the center of targeted trees. b. and d. perspective scene view in Google Earth, with yellow pins indicating camera locations along each trajectory at the moment images were captured.

**Table 2.** Mission Planner commands used for autonomous UAV flights

| Command | Code | Description |
| --- | --- | --- |
| WAYPOINT | 16 | Latitude, longitude (in degrees) and altitude vector (in meters) of locations visited during a flight |
| DO_CHANGE_SPEED | 178 | Speed, in meters per second. Calculated considering distance to target and image acquisition frequency, usually 2Hz. Typical speed value is 4 meters per second |
| DO_SET_ROI | 201 | Vector of UAV heading planar azimuth and gimbal angle (in degrees) that orients the camera towards relative to a specified point of interest. |
| RETURN_TO_LAUNCH | 20 | Return to launch location after flight completion |
| DO_SET_HOME | 179 | Latitude and longitude vector (in degrees) of return UAV location to use in the event of an emergency, or system anomaly |

### 2.4 Evaluation of tree reconstructions

Processing of the synthetic imagery always produced complete tree reconstructions. The number of points in the reconstruction ranged between 20 and 25 percent of those present in the original terrestrial LiDAR point cloud (Figure 3b). Larger percentages could be achieved by increasing the resolution of the virtual camera, at the expense of prolonged processing time in both VSfM and SURE. Volumetric comparisons in voxel space revealed excellent agreement between LiDAR and reconstructed point clouds, with a mean of 94 percent of filled voxels collocated. Omnidirectional jittering of the voxel-rendered tree reconstructions relative to the terrestrial LiDAR equivalent always resulted in a substantial, 30 to 40 percent reduction in collocation rates, even when the jittering was limited to a single voxel. The rapid reduction in the collocation rates caused by jittering limited to one voxel suggests that the scaling and translation of the derived point cloud relative to the original terrestrial LiDAR cloud is accurate and precise. It also implies that the coordinates of the virtual camera positions deduced by VSfM during the processing of the synthetic imagery and those used in the simulation are identical up to the scale difference. Once calculated, scaling and translation of the reconstructed point cloud performed by using this relationship rendered the derived tree point cloud a thinned copy of the original terrestrial point cloud. Our simulation results suggest that the absence of planar surfaces and lack of opacity in tree crowns do not impose systemic restrictions to the surface-from-motion approach we used to obtain the 3D tree representations.

By exploring several virtual camera trajectory patterns while altering the image acquisition frequency in each of them, we were able to quantify the effects that different patterns and image field-of-view overlap percentages have on tree reconstruction accuracy (Figure 7). Even in the ideal, noise-free environment of the simulations, a minimum 30 percent image overlap was required for complete target reconstructions. For patterns involving camera locations at variable above-ground heights the minimum percentage was higher, between 35 and 40 percent. Below a mean 45 percent overlap, all simulations were susceptible to failure, pending on the image pair selected for initiating the matching process described in section 2.1. For the circular trajectory pattern, the level of volumetric correspondence between the terrestrial LiDAR and imagery-derived point clouds would increase rapidly at low field-of-view overlap percentages and then progressively decline until reaching an asymptote, usually at 90 percent volumetric correspondence or higher (Figure 7). Complete reconstructions obtained with the spiral trajectory usually required at least 35 percent image overlap. The observed volumetric correspondence to the LiDAR point cloud showed a sigmoidal increase with higher image overlap percentages until reaching an asymptote level, sometimes as high as 94 percent.

**Figure 7.**
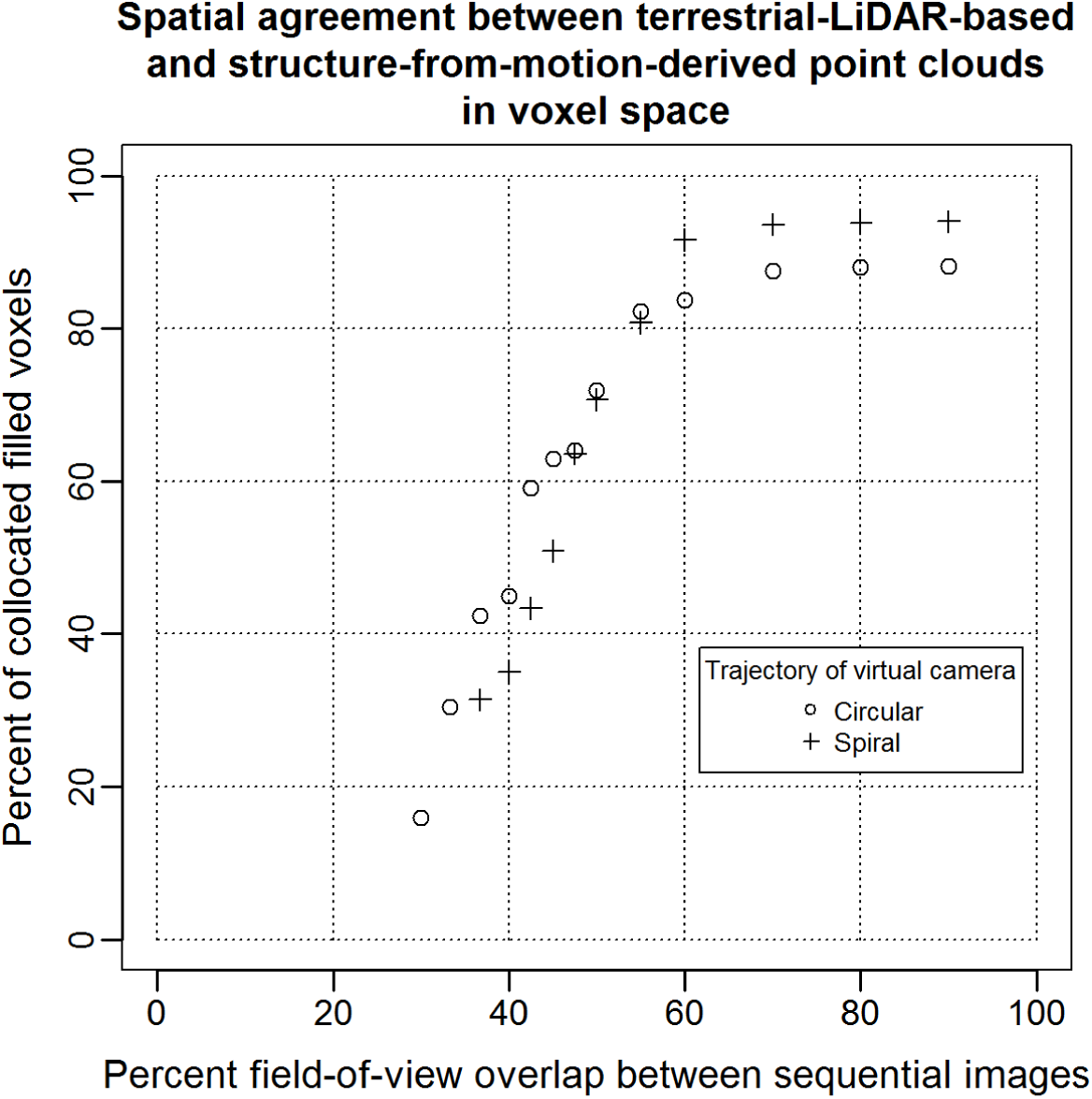
Accuracy and completeness of reconstruction for a *Pinus ponderosa* tree. This analysis is based on synthetic imagery simulated using visualization of terrestrial LiDAR point clouds and two camera trajectories. Percentage of collocated filled voxels is used as reconstruction completeness criterion.

In a spiral acquisition trajectory yielding the same number of images of a targeted tree as a circular trajectory, the horizontal overlap percentage between two sequential images is lower. Unlike the circular trajectory, though, in the spiral there is vertical overlap with images obtained after the UAV has completed a rotation around the tree. While the overall mean overlap between the two trajectory patterns was the same in our simulations, the spiral had lower overlap percentage between any two images selected for the initiation of the matching process, and therefore more likely to fail to yield a complete reconstruction when the overall overlap image rate was low. Owing to the vertical image overlap present in spiral UAV missions, selected parts of the tree are visible from more than one vertical viewing angles, an arrangement that reduces target occlusion rates. For tree species with dense, uniform distribution of foliage and deeply shaded crown centers, the variability in vertical view angles offered by the spiral trajectory pattern may be unimportant. For species with predominantly horizontal or angular branch arrangement and lower crown compaction rates, vertical viewing variability allows internal crown components to be represented adequately in the derived point cloud. Three out of four of the voxels accounting for the approximately 4 percent difference in reconstruction completeness between the spiral and circular UAV trajectories around a Red Pine (*Pinus ponderosa*) tree at 70 percent image overlap rates or higher (Figure 7) were located in the internal half of the crown.

Most UAV flights also produced complete tree reconstructions (Figures 8 and 9). In the absence of detailed crown dimensionality measurements, we relied on ocular assessment of reconstruction accuracy and precision. The typical example shown on Figure 8, obtained with the spiral UAV trajectory (Figure 5c), among our most reliable for complete target reconstruction, shows that even the shaded components of the tree crown interior are represented. Many parts on the upper quarter of the crown have a light blue hue inherited from the sky background in corresponding UAV images. Although less evident, selected parts of the lower crown exhibit similar ground-influenced coloring. The coloring artifacts shown in Figure 8 appear where the image area occupied by an identified keypoint is dominated by a uniformly colored background. Sometimes these anomalies are limited to the RGB values assigned to points and can be overlooked if the main objective of the UAV mission is the retrieval of tree dimensionality. Often though they represent an overestimation of tree crown volume and must be removed (Figure 10). Accomplishing this task with manual intervention is laborious and subjective. The task can be easily automated for points pertaining to a sky background thanks to their markedly different RGB values compared to those of vegetation. Where suitable RGB value thresholds cannot be safely identified, as it is usually the case for the lower parts of trees, we found it useful to trim the depth of the part of the overall reconstructions that is derived from each image, so that only the portion nearer the camera position is retained. SURE facilitates this procedure by providing a separate dense reconstruction for each processed image organized in a common coordinate system. The complete reconstruction can be obtained by merging the trimmed parts. In the absence of precise reference data, we were unable to determine quantitatively the significance of these artifacts.

**Figure 8.**
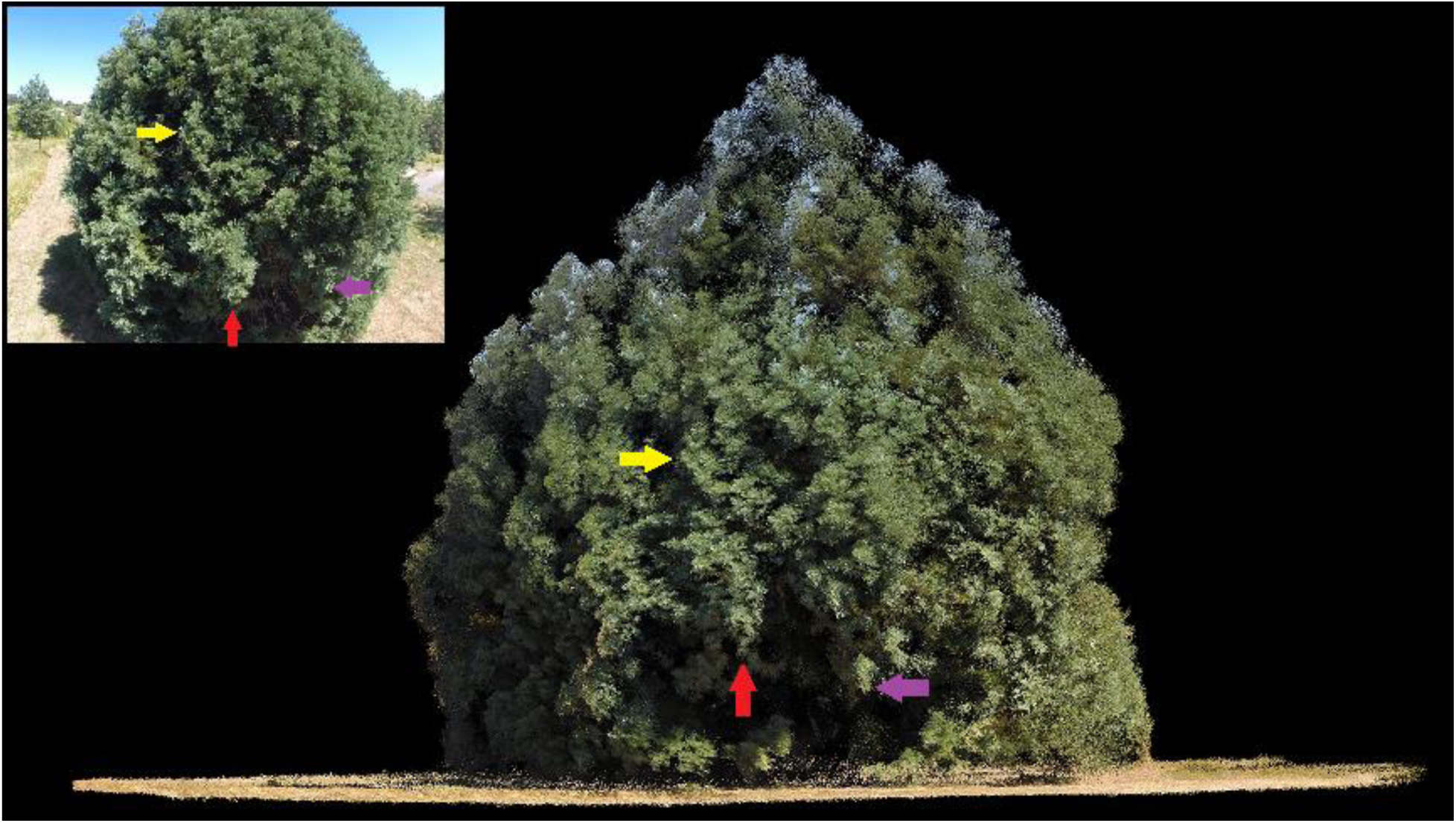
Orthographic horizontal view of reconstructed point cloud and UAV-based oblique perspective image. Colored arrows denote corresponding tree crown components.

**Figure 9.**
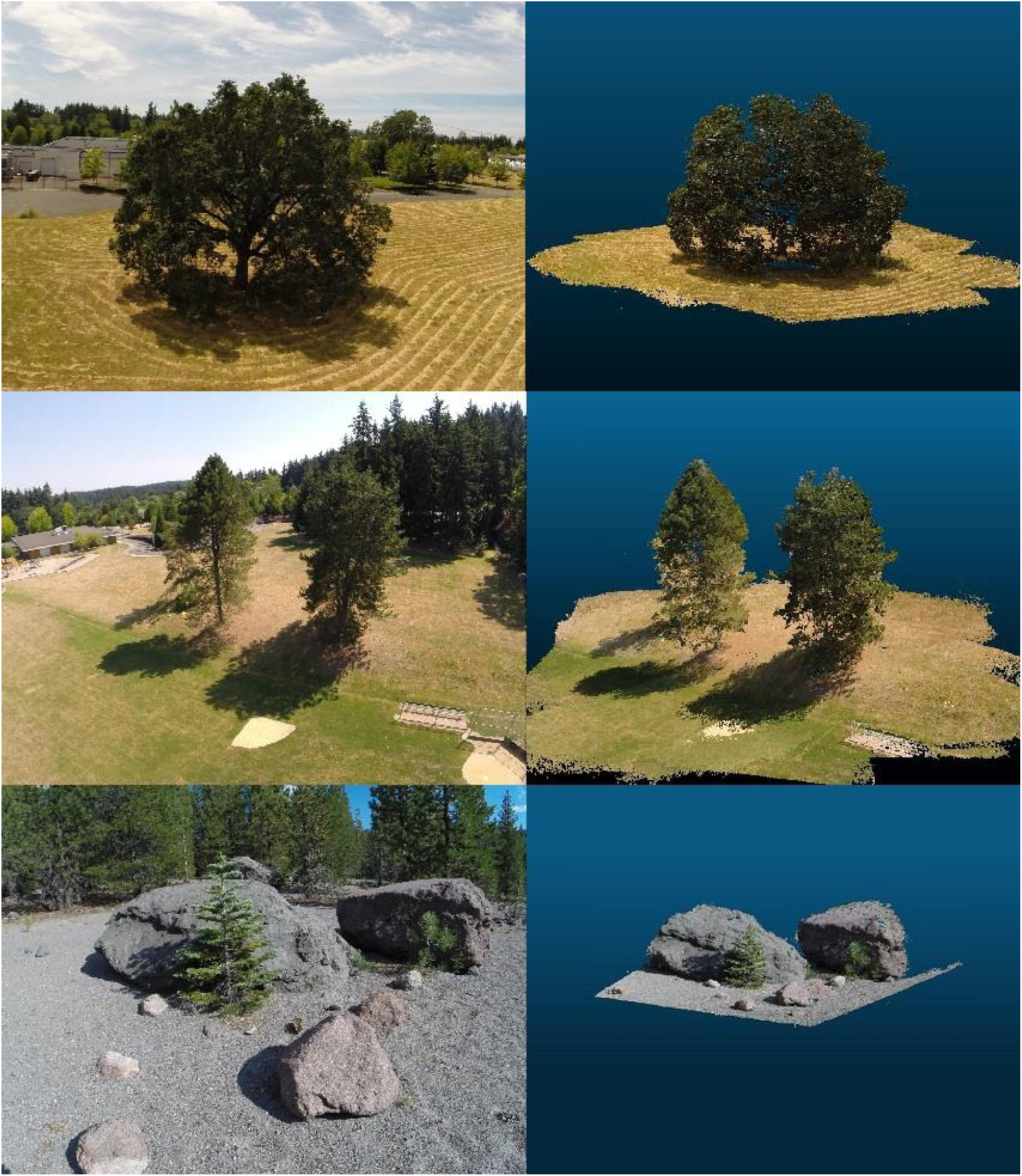
Illustration of comprehensive tree reconstructions (right column) and reference UAV-based images (left column).

**Figure 10.**
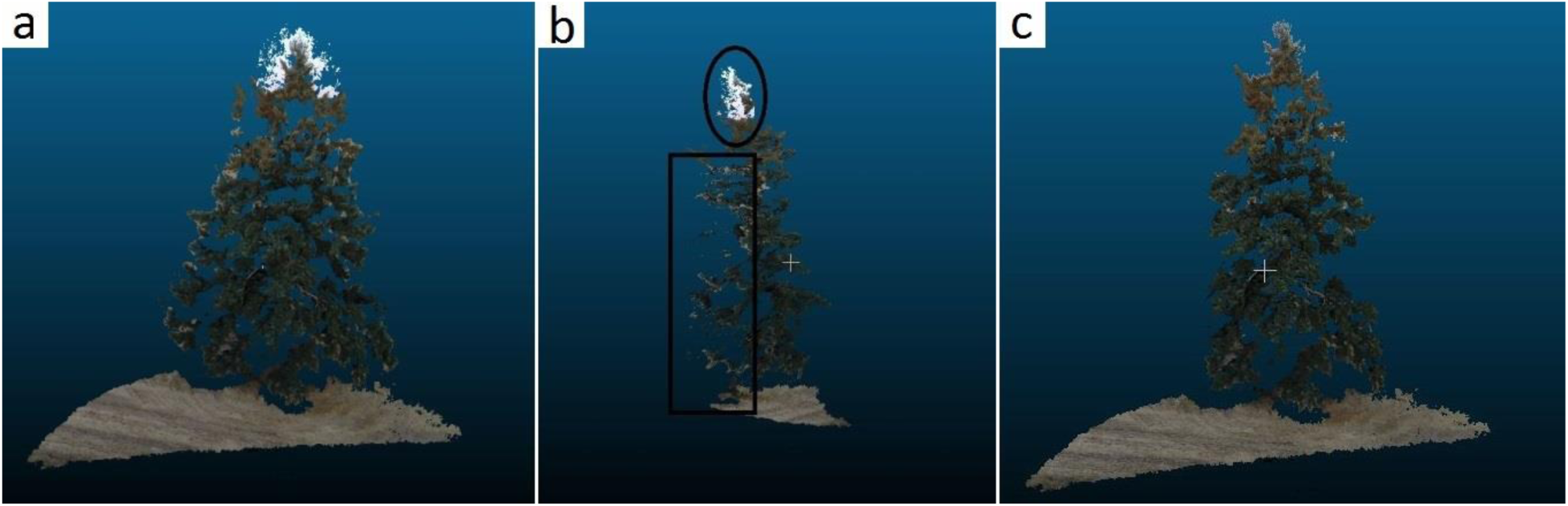
Demonstration of artifacts in the 3D tree reconstruction pertaining to a single UAV image. a. Initial reconstruction, positioned facing the camera with a band of white-colored points belonging to sky background near the top, and light colored points to the sides belonging to fallow land background, b. Side view, with camera position to the left and sky points in oval and land points in rectangle, and c. Trimmed reconstruction positioned facing the camera.

The ‘nested circle’ and ‘jagged saucer’ trajectories (Figure 5g and 5h) produced only partial reconstructions and several disjointed models in VSfM and are, therefore, not recommended, while the altitude variability in the ‘meandering’ trajectory (Figure 5d) was often responsible for premature mission termination owing to rapid depletion of the UAV batteries. Partial reconstructions were the norm, rather than the exception, when for a portion of the mission the camera was positioned directly against the sun. In such conditions the shaded portion of the crown would either not be reconstructed at all, or it would be organized in separate 3D models with much lower point density and sizable gaps. In the example shown in Figure 11, the GPS recorded and process-derived positions of the camera on board the UAV show a nearly perfect correspondence for three quarters of the circular UAV trajectory. GPS recordings are half as many as the camera positions because of limitations in the recording frequency of the GPS device. Is should be noted that pending on the hardware configuration of the UAV and the number of peripheral devices connected to it, it is sometimes necessary to operate below the capacity of a particular device to either conserve energy, or to avoid overwhelming the UAV controller. Based on our experience, a close fit between recorded and derived camera positions practically guarantees that a complete target representation will be obtained during the dense reconstruction phase. The remaining part of the trajectory, where the camera is positioned against the sun, was actually derived from a separate model and shows a poor fit, resembling more of a linear transect than a circular arc. As the camera moves from partially to completely against the sun, image contrast is reduced, and the radii of identified keypoints become smaller. Radius reductions increase the uncertainty associated with keypoints orientation and descriptor. We suspect that changes in the magnitude of the mean image keypoint radius are manifested as variability in the distance between the tree and calculated camera locations, evident in the misfit part of the VSfM-derived camera trajectory shown in Figure 11.

**Figure 11.**
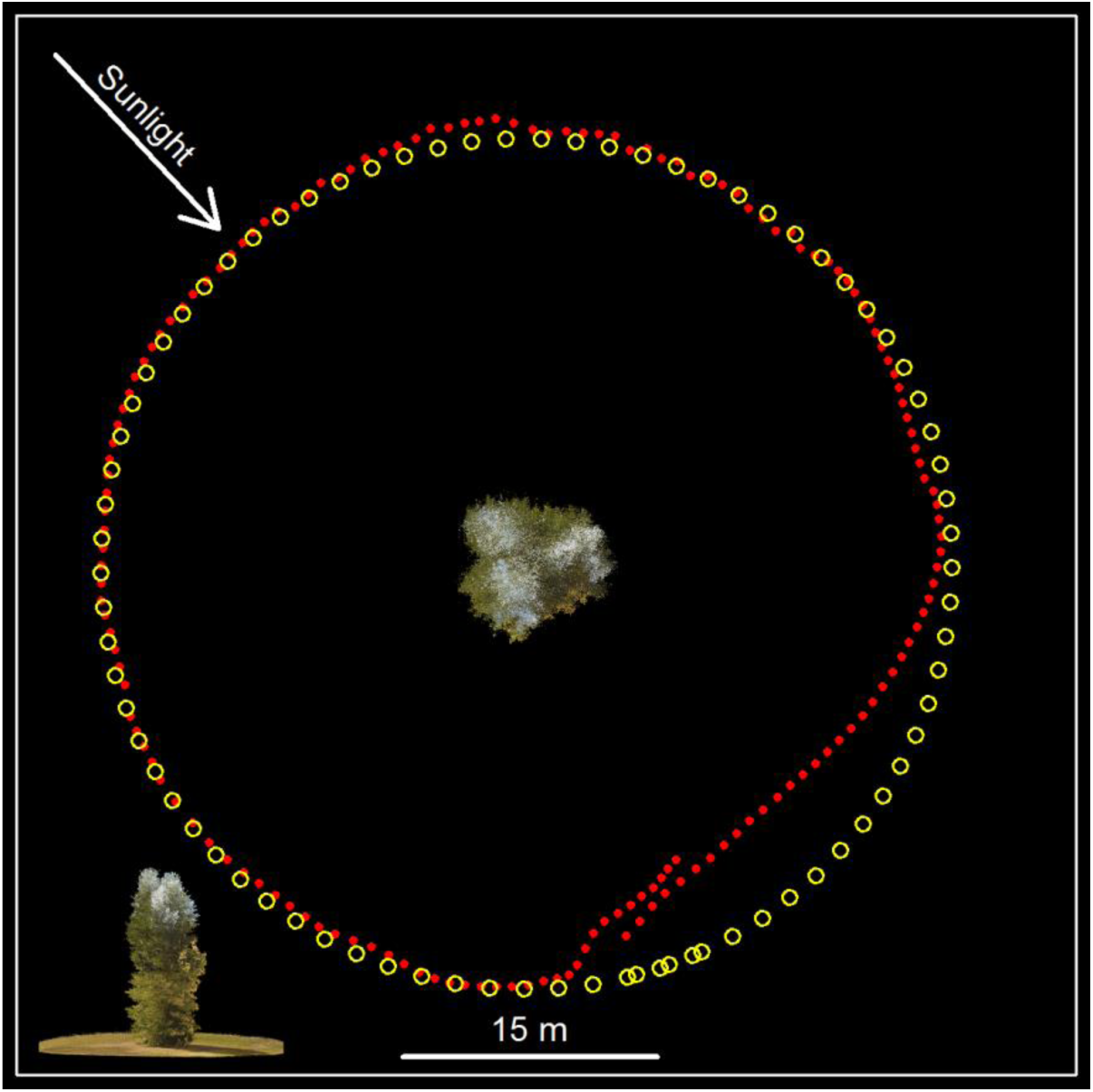
Comparison between real and reconstructed trajectory. Nadir view of reconstructed tree with camera GPS locations at image frame acquisition moments (yellow circles) and VSfM-calculated locations (red dots). Frame frequency 2Hz, GPS fixes at 1Hz. Inset at the lower left shows lateral view of the reconstructed tree.

On a few occasions, we observed more than one, nearly parallel, and closely stacked layers of points representing the ground, likely an artifact of texture uniformity in those parts of the scene. The use of calibrated imagery has expedited the computations for identifying camera positions and for generating the sparse reconstructions in VSfM and has reduced the rate of partial reconstruction occurrence. However, its effect on the accuracy of the reconstruction obtained using SURE was unclear.

## 3. Discussion

Our results indicate that a meticulously planned image acquisition mission, namely a judicious selection of flight trajectory, UAV speed, and image acquisition frequency, will deliver a comprehensive dense reconstruction of targeted vegetation, except perhaps in unfavorable sun illumination and wind conditions. As explained in section 2.1, our workflow relies on keypoints, most of which are identified along image discontinuities. A smooth flight trajectory around the target ensures that sequential images contain an adequate number of similar keypoints from which the camera location effective for each image capture can be calculated with adequate precision. Where the smooth change in the field of view between two sequential images is interrupted, the offending image becomes the first in a separate model. Bundle adjustments can reduce the frequency of separate model emergence but they cannot eliminate it. The often advocated practice of adding to a model image frames originally put by VSfM to a separate model without performing bundle adjustment after each frame addition may be warranted for manmade objects but is not recommended for trees because it leads to obvious reconstruction artifacts. Mission plans for flights expected to occur during bright solar illumination conditions using gimbal-equipped UAVs could be adjusted to avoid camera positioning directly against the sun. This can be accomplished by specifying a slightly downward, oblique camera orientation. The precise solar elevation angle and azimuth for any location can be obtained from the NOAA Solar Position Calculator (http://www.esrl.noaa.gov/gmd/grad/solcalc/azel.html), or can be computed as described in Reda & Andreas (2008).

GPS-equipped UAV platforms not only enable preprogrammed navigation, but also, and perhaps equally importantly, can be used for a precise scaling of reconstructed tree point clouds to actual dimensions. The GPS receivers placed on the two UAVs employed in this study offer recreational grade precision, and as such, their individual position recordings may contain an absolute error of a few meters. In our trials, however, the relative error between trajectory recordings appeared to always be less than a meter, in most cases about half a meter. This is based on the observation that our UAVs, initially placed on a launch pad measuring about 60 cm on each side, would return at the completion of the mission with their landing gear partially on the launch pad. Fitting the VSfM-calculated camera locations to corresponding GPS recordings containing a relative positional error of such magnitude, would yield point cloud scaling errors of 0.5 percent or lower, a level deemed adequate for UAV imagery and structure from motion based assessment of yearly tree growth. In the absence of GPS recordings, the scaling of the point cloud would have to be performed manually using georeferenced imagery.

Except for extremes in solar illumination conditions such as sun facing camera exposures or at dusk, disparities in light distribution may actually be beneficial for structure-from-motion-based applications in natural environments because they accentuate feature edges. As it is evident in the tree portion between the red and purple colored arrows shown in Figure 8, crown parts in the penumbra are still represented, albeit with reduced point density. Image enhancements focusing on shaded or very bright parts could perhaps be used to ameliorate the direct sunlight effects or improve the reconstruction density for shaded areas.

To account for absolute GPS receiver and ancillary imagery registration errors, current UAV missions must be planned with adequate clearance from any scene objects. We were able to comply with this requirement in our trials because we mostly targeted individual trees or small groups of trees growing in open space. Extending our operations to confined areas, for instance descending into and proceeding near and along the periphery of forest openings, would require much higher navigation precision. Thankfully, obstacle avoidance has been actively researched and several solutions specific to forested environments have been proposed (Frew et al., 2006; Karaman and Frazzoli, 2012; Mori and Scherer, 2013; Roberts et al., 2012; Ross et al., 2013). In particular, Ross et al. (2013) demonstrated full flight control in forested environments using an UAV platform similar to ours. They used a low-resolution camera mounted on a quadcopter that was outsourcing via a wireless connection all computationally intensive image processing to a ground station, a standard laptop computer. Using this setup, they were able to achieve a constant speed of 1.5 meters per second while avoiding trees. The rapidly expanding onboard processing capabilities of UAVs suggest the possibility, in the near future, of coupling the 3D reconstruction methodology proposed here with autonomous flight, thereby eliminating the need for meticulous mission planning.

It is often tempting to acquire images with the highest possible frequency and maximum overlap. Action cameras similar to those used in this study support high frame rates and carry ample image storage space without affecting the payload and thus compromising the UAV’s flight duration or mission flexibility anyway. Large number of images though requires prolonged processing. Our simulations indicate that image field-of-view overlap higher than 70 percent, does not improve the accuracy or completeness of tree reconstructions. Visual assessments suggest that this is also true for actual UAV imagery. Mission planning designed so that target features are represented in three to four images likely maximizes the information content present in an acquisition and it is therefore recommended as an initial mission configuration.

## 4. Conclusion

Rapid developments in UAV technology and enhancements in structure from motion software have enabled detailed representation of manmade objects. In this paper, we describe how this technology can inexpensively be extended to representations of natural objects, such as trees or groups of trees. After extensive experimentation that involved several UAV platforms, cameras, mission planning alternatives, processing software, and numerous procedural modifications and adjustments, our workflow has been proven capable of handling most conditions encountered in practice to deliver detailed reconstruction of trees. In addition to robust performance, our imaging system can be employed rapidly in support of time-sensitive monitoring operations as, for instance, the assessment of forest fire damage or progress of forest recovery from disturbance. It is also well suited to providing tree dimensionality data through time, a prerequisite for improved models of tree growth and for an accurate assessment of tree competition and morphological plasticity.

